# Assessment of the normal cell contamination impact on tumour sample analysed with SNP arrays: The signal confusion nightmare

**DOI:** 10.1101/2023.04.04.534870

**Authors:** Christophe B. Poulet, James T. Swingland, Vincent Botta, Pierre Robe, Christian Herens, Federico Turkheimer, Vincent Bours

## Abstract

Recent advances in high-throughput technologies enable a more comprehensive interpretation of the tumour evolution through the study of the intra-tumour heterogeneity. Several algorithms, however, often relies on the use of models that described the top of the iceberg regarding the stromal contamination of the samples, making diagnosis difficult to assess. Indeed, such as radio wave receivers, tools to analyse high-throughput technologies data, are used to enable the discrimination between multiple signals differing in frequencies. However, such tools often look at the average frequency more than distinct signals, leading to analyse a confused signal. This confusion could dramatically lead to a mis–interpretation of the real data, especially during the diagnosis as it relies on the choice of a unique scenario among many others. Here, we describe how this signal confusion occurs in the most classical DNA microarray analysis of tumours and we provide statistics to determine how many other possible scenario can lead the same signals, in order to improve the robustness of pigeon hole logic based analysis. Based on simulations, where a unique tumour population was diluted by an increasing gradient of normal cells, we underline the causes and consequences of such signal confusion for up to five allelic copies. Despite the removal of all technical biaises and background noise, we show how the signal confusion remains systematically present in the commonly used DNA microarray analysis, especially for the genotypes AAAAB, AAAB and AAB for copy numbers 5, 4 and 3 respectively, as well as their symmetric combinations for the B allele.

## Introduction

Cancer genomes are often distinguishable from normal genomes by their numerous genetic aberrations. These aberrations can take several forms from punctual mutations to chromosomal rearrangements such as gains, losses, translocations or inversions. Many of these events were linked to specific tumour types, and now constitute strong markers for the cancer characterisation and classification [1, 2]. These events were acquired and selected during the long process of tumourigenesis. Indeed, tumours are the result of a long evolution that includes multiple rounds of proliferation, cell death, and genomic rearrangements. This process implies a continuous genesis of new aberrations that are not always ubiquitous through tumour cells [1-8]. The most striking events are gains and losses, which are commonly called copy number alterations (CNA). These CNAs often produce allelic imbalances (Ai), which are an over-representation of one of the two original heterozygous alleles. The most striking Ai is the loss of heterozygosity (LOH), where a normal heterozygous locus become homozygous after a loss event. All non LOH allelic imbalances, are called here other Ai (oAi).

Single nucleotide polymorphism (SNP) arrays (SNP–aCGH) allow the screening of CNAs and Ais through the allele specific analysis. These technologies measure the intensity of fragmented genomic DNA sequences that are hybridised to probes which cover most part of the genome. The allele specific analysis is enabled by the targeted sequences. Indeed, probes target genomic regions of known SNPs and thus measure the two possible allelic versions of this SNP. For convenience, these two possible versions are labelled A and B alleles. Thus, these platforms provide both the Log R ratio (LRR), the total intensity of both A and B alleles, as well as the B allele frequency (BAF), the relative proportion of the B allele. While the reproducibility of these platforms is not to debate here, the biopsy composition can bias the resulting signal.

Along the tumourigenesis, cell clones are selected for their survival and proliferation advantages. Although most cancer cells are expected to share common genetic features, coming from their most recent common ancestor, the recent advances in high-throughput genetics emphasised significant sub–populations in most tumours [8, 11, 12]. Even if a sub–population exhibits a high fitness, it rarely reached the fixation at the time of the surgery, thus resulting in the co-existence of several sub–populations in the sample [3, 7, 13]. Consequently, tumours are often composed of a mixture of genetically different cell sub-populations which constitutes the *intra*-tumour heterogeneity (ITH). Like waves in physics, the detected signals are the subject of interferences, such as signal destruction, which often lead to a mis–interpretation of the data. This signal destruction can occur for LRR and BAF signals when two equally represented populations carry opposite aberrations, such as gain and loss with complementary genotypes, such as A and ABB.

Tumour samples picked-up by surgery are often contaminated with normal stromal cells. This source of additional interferences makes the overall sensitivity of LRR and BAF signals proportionally distorted along with an increasing amount of normal cells. Such contamination often led to difficulties in the assessment the exact CNA or the detection of LOH events [14]. Furthermore, stromal cells were previously described as an additional source of chromosomal aberrations, making the overall mixture even more complex [15, 16].

Tools to address all these sample effects were developed during the last decade [4, 14, 17-22]. The main attempt of these algorithms is to assess the exact copy-number status of such complex samples. For this purpose, the LRR and the BAF signals were combined to circumvent the signal distortion. Two distinct methods were mainly used, hidden Markov models (HMM) and change point detection (CPD). Studies that compared these tools concluded that CPD based algorithms outperform HMMs [22]. Besides, a pigeon hole logic can be applied to the entire set of individual alterations to improve the exact CNA assessment. Altogether, these tools were able to solve most of the crucial SNP–aCGH problems, such as noise, polyploidy as well as ITH.

However, the complex genetic of the tumour samples remains difficult to assess. A high number of tumour sub–populations with different genetics often led to a ill-mathematical problem during the assessment of the exact CN status [4]. This problem, called here “*Signal Confusion*”, is one of the basic failures encountered by SNP– aCGH analysis, and we expect a comparable confusion to occur only with a stromal contamination. Indeed, current tools often consider the stromal contamination as an additional population, which only lead to a signal reduction, where the LRR and BAF signals converge to the neutral signal of a diploid state [4, 9]. Moreover, it is not known which genotypes are more likely to be the source of such signal confusion. Finally, even though efficients, pigeon hole algorithms have to choose among numerous equally probable scenario of combined individual alterations. Consequently, SNP-aCGH analysis may not always reflect the exact reality, and thus, may increases the number of analysis biases, leading to a quick interpretation and thus a false conclusion during diagnosis.

Here, we assessed the impact of such stromal contamination on the common SNP–aCGH analysis, with the aim to identify the basic sources of signal confusion occurrences. Furthermore, we developed statistics to weight each signal for any confusion occurrence. These probabilities can be used as a degree of certitude on each individual alteration, but also to assess the amount of equally probable scenario that can result from a pigeon hole analysis. To keep focus on the expected effect of signal confusion, we avoided any additional background noise, polyploidy or other ITH in the analysis. Thus, we analysed optimised simulated signals from a population that carry CNs from *CN*_0_ (bi-allelic deletion) to *CN*_5_ (amplification), diluted in an increasing gradient of 1−α% of normal cells (*CN*_2_) from 0 to 100%.

## Results

### Copy Number analysis

To assess the impact of normal cells on the exact copy number (CN) analysis, we looked for the signal confusion on the Log R ratio (LRR). For this purpose, we simulated LRR signals for CNs from *CN*_0_ to *CN*_5_ using SiDCoN simulation algorithm [23]. Each signal was simulated with an increasing gradient of tumour cells (α) from 0% to 100%. Figure 1 summarises the evolution of the LRR signal according to the α% of altered cells in the sample, for each simulated CN. With the exception of the *CN*_0_, each CN describes a straight line of LRR signal function of α% of altered cells. Importantly, several signals can be achieved with different combination of CN and α. As exemplified by the dashed lines and listed in table 1, the signal *LRR*=−0.415 can be achieved both by a *CN*_1_ at α=76% of altered cells and a *CN*_0_ at α=38%. Similarly, the signal *LRR*=0.323 can be achieved either by a *CN*_3_ at α=90%, a *CN*_4_ at α=45% or a *CN*_5_ at α=30%. Every genotype combinations that converged to the same CN ended with the same LRR signal at the same α. The α% of altered cells can be found based on the LRR signal and the CN status with the equation 6 (see Methods), adapted from SiDCoN.

**Figure 1.**
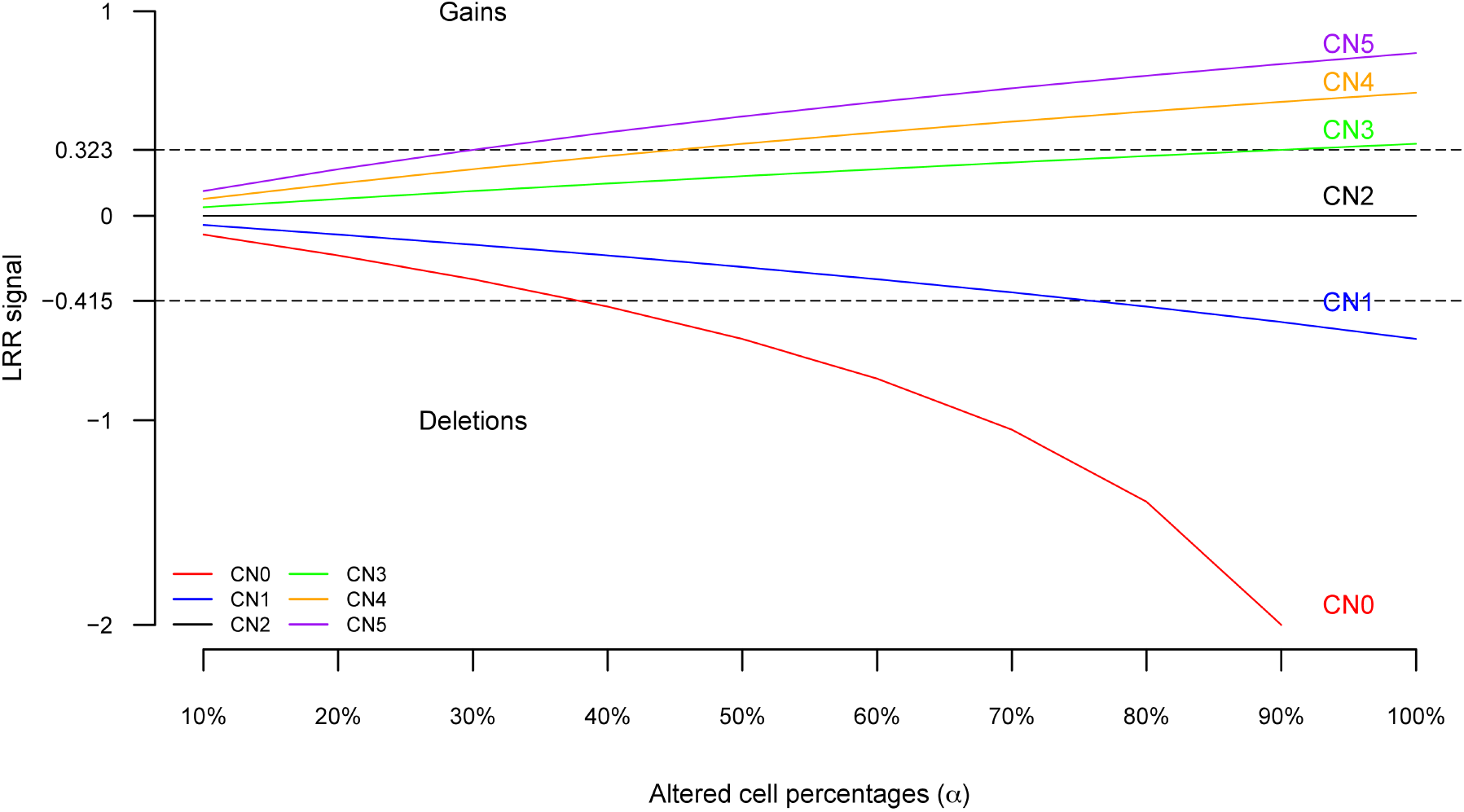
Signal Confusion in Copy Number analysis. Evolution of the LRR signal against α% of tumour cells that carry CNs from 0 to 5 (red to purple). Dashed lines highlight signal confusion for -0.415 and 0.323 LRR signals. Each value of the graph was simulated based on SiDCoN algorithm. Except for the bi-allelic deletion (*CN*_0_), a copy number defines a straight line according to the altered cell percentage. Most of LRR signals can be achieved with different combination of CNs and α% of altered cells.

**Table 1:** LRR’s signal confusion example.

| **LRR** | **CN** | $N_A^t$ | $N_B^t$ | $\alpha$ |
| --- | --- | --- | --- | --- |
| -0.415 | 0 | 0 | 0 | 0.38 |
| -0.415 | 1 | 0 | 1 | 0.76 |
| -0.415 | 1 | 1 | 0 | 0.76 |
| 0.323 | 3 | 0 | 3 | 0.9 |
| 0.323 | 3 | 1 | 2 | 0.9 |
| 0.323 | 3 | 2 | 1 | 0.9 |
| 0.323 | 3 | 3 | 0 | 0.9 |
| 0.323 | 4 | 0 | 4 | 0.45 |
| 0.323 | 4 | 1 | 3 | 0.45 |
| 0.323 | 4 | 2 | 2 | 0.45 |
| 0.323 | 4 | 3 | 1 | 0.45 |
| 0.323 | 4 | 4 | 0 | 0.45 |
| 0.323 | 5 | 0 | 5 | 0.3 |
| 0.323 | 5 | 1 | 4 | 0.3 |
| 0.323 | 5 | 2 | 3 | 0.3 |
| 0.323 | 5 | 3 | 2 | 0.3 |
| 0.323 | 5 | 4 | 1 | 0.3 |
| 0.323 | 5 | 5 | 0 | 0.3 |

**LRR**: Observed Log R ratio, **CN**: Tumour's Copy Number, $N_A^t$ : Tumour's A allele amount, $N_B^t$ : Tumour's B allele amount, $\alpha$ : % of tumour cells.

### Allelic analysis

To assess the impact of normal cells on the allelic imbalance (Ai) analysis, we looked for the signal confusion on the Θ and the B allele frequency (BAF) analysis, as well as any disturbance that could occur in the LOH detection analysis. For this purpose, we simulated the Θ and BAF signals for all allelic combinations (AA, ABB, *etc*.) that led to tumour CNs from *CN*_0_ to *CN*_5_. Each signal was simulated with an increasing gradient of tumour cells (α) from 0% to 100%. Interestingly, all aberrations signals that derived from the homozygous genotypes (AA or BB) led to the same signal, which increased the signal confusion. Thus, only the aberrations that came from the heterozygous genotype (AB) were considered here.

#### Theta (Θ)

Figure 2 summarises the different combinations of Θ values obtained for the dilutions of altered cells with CNs from *CN*_0_ to *CN*_5_. Θ values were calculated based on Wang *et al* [24]. The signals varied between 0 and 1, and were symmetric against 0.5. They also described three distinct curve shapes, that depended on the allelic combination. Balanced alleles combination (triangles), described a straight line at 0.5 (figure 2.b). They occurred for even CNs when alleles were equally represented. LOH (circles), described curves function of α (figure 2.c). They occurred when only one allele was represented. Their signals started from Θ=0.5 to one of the two extrema, depending on the allele involved (A or B). Other Ais (squares), described also curves function of α (figure 2.d). They occurred when one allele is over–represented. These signals also started from Θ=0.5, but ended before the maxima, 0<Θ<1, depending on the most represented allele. It is worth to note that many Θ signals can be achieved with different CN or combinations of alleles at different α. As exemplified by the dashed line and reported in the table 2, a Θ signal at 0.3 can be achieved by eight different combinations of genotype and α. Additionally, all aberrant genotypes coming from a AA reference led to a Θ value of 0. Similarly, the aberrant genotypes coming from a BB reference led to a Θ value of 1. Any α can be found based on the Θ signal and the amount ***N***_***A***_^**t**^and ***N***_***B***_^**t**^of allele A and B respectively, with the equation 7 (see Methods), adapted from Wang’s equation [24].

**Figure 2.**
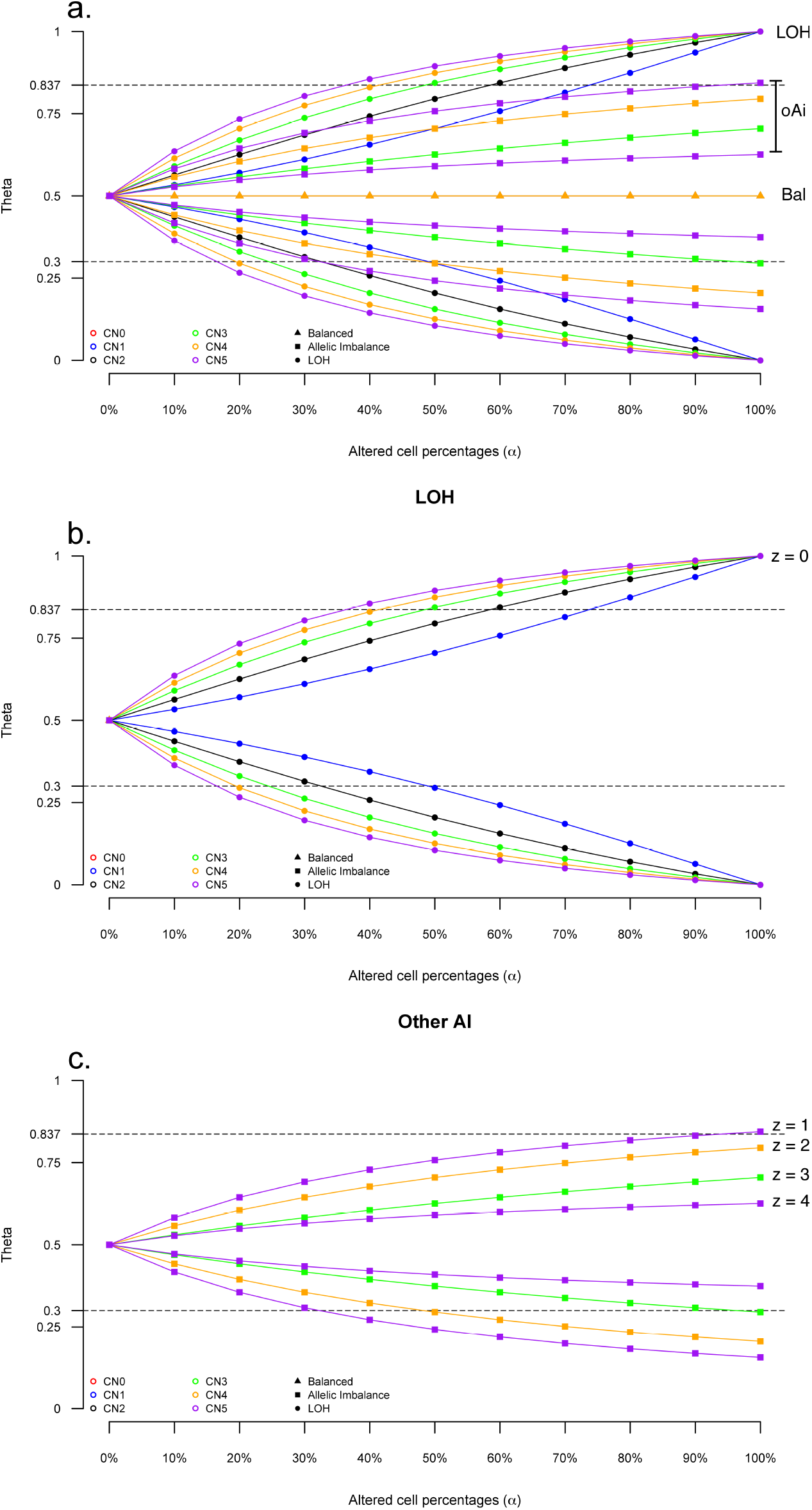
Signal Confusion in Theta (Θ) analysis. Evolution of the Θ signal against α% for CNs from 0 to 5 (red to purple). Each value of the graph was simulated using Wang’s method to estimate Θ. **a**. All possible genotypes that led to CNs from 0 to 5 which can be separated into three genotype classes. Balanced (triangles). LOH (circles). Other Ais (squares). **b**. LOH genotypes only (circles). **c**. Other Ai genotypes only (squares). Dashed lines highlight signal confusion for Θ=0.3 and Θ=0.837. Every Θ signals can be achieved with different allelic combinations and α% of altered cells.

**Table 2:** Θ’s signal confusion example.

| $\Theta$ | CN | $N_A^t$ | $N_B^t$ | $\alpha$ |
| --- | --- | --- | --- | --- |
| 0.3 | 1 | 1 | 0 | 0.49 |
| 0.3 | 2 | 2 | 0 | 0.32 |
| 0.3 | 3 | 2 | 1 | 0.96 |
| 0.3 | 3 | 3 | 0 | 0.24 |
| 0.3 | 4 | 3 | 1 | 0.48 |
| 0.3 | 4 | 4 | 0 | 0.19 |
| 0.3 | 5 | 4 | 1 | 0.32 |
| 0.3 | 5 | 5 | 0 | 0.16 |

$\Theta$ : Observed $\Theta$ signal, CN: Tumour's Copy Number, $N_A^t$ : Tumour's A allele amount, $N_B^t$ : Tumour's B allele amount, $\alpha$ : % of tumour cells.

#### BAF

Two different methods were used to calculate the BAF in previous studies, the Θ based BAF, derived from Wang’s method [24] and the classical BAF, derived from SiDCoN algorithm [23]. Both methods were compared for their behaviour against a stromal contamination. For this purpose, we measured the signal confusion on the mirrored BAF (mBAF), which constrains the signal between 0 and 0.5. Figure 3.a summarises the different combinations of mBAF values, obtained using Wang *et al* algorithm [24]. These mBAF signals were obtained for CNs from *CN*_0_ to *CN*_5_, diluted with an increasing amount of normal cells. Similarly, figure 3.b summarises the same signal of mBAF, but estimated with the SiDCoN algorithm [23]. With the exception of balanced allelic amounts (triangles), such as the *CN*_0_, *CN*_2_ and *CN*_4_, each allelic combination had a mBAF signal over 0. Like the Θ signal, the mBAF signal described two other types of curves, depending on the allelic combination. LOH (circles), described curves function of α linking 0 to the maximum of 0.5. Other Ais (squares), described curves function of α, linking 0 to a maximum value below 0.5. This maximum value followed the disequilibrium between the two alleles in the genotype. Importantly, in both methods, almost all mBAF signals can be achieved with different combinations of alleles for different CNs and α. As exemplified by the dashed line in figure 3 (a) and reported in the table 3, the Θ based mBAF signal at 0.3, estimated using Wang’s method, can be achieved with all combination of LOH or two other oAi genotypes and α. Similarly, the dashed line in figure 3 (b) showed a classical mBAF signal at 0.3, estimated with SiDCoN’s method. As reported in the table 4, this mBAF signal can be achieved with all combination of LOH genotypes and α, but only one oAi and α combination. Moreover, the Θ based mBAF at 0.5 can be achieved with all LOH combinations indifferently for α≥0.5. For the Θ based mBAF, any α can be estimated with equation 8 (see Methods), which was derived from a Wang’s algorithm. For the classical based mBAF, any α can be estimated with the equation 9 (see Methods), which was derived from SiDCoN method. Interestingly, in both BAF cases, all aberrant genotypes derived from a AA or a BB reference led to a mBAF value of 0.5.

**Table 3:** Θ BAF signal confusion example.

| **mBAF** | **BAF** | **CN** | $N_A^t$ | $N_B^t$ | $\alpha$ |
| --- | --- | --- | --- | --- | --- |
| 0.3 | 0.2 | 1 | 1 | 0 | 0.57 |
| 0.3 | 0.2 | 2 | 2 | 0 | 0.4 |
| 0.3 | 0.2 | 3 | 3 | 0 | 0.3 |
| 0.3 | 0.2 | 4 | 3 | 1 | 0.66 |
| 0.3 | 0.2 | 4 | 4 | 0 | 0.25 |
| 0.3 | 0.2 | 5 | 4 | 1 | 0.44 |
| 0.3 | 0.2 | 5 | 5 | 0 | 0.21 |
| 0.3 | 0.8 | 1 | 0 | 1 | 0.57 |
| 0.3 | 0.8 | 2 | 0 | 2 | 0.4 |
| 0.3 | 0.8 | 3 | 0 | 3 | 0.3 |
| 0.3 | 0.8 | 4 | 0 | 4 | 0.25 |
| 0.3 | 0.8 | 4 | 1 | 3 | 0.66 |
| 0.3 | 0.8 | 5 | 0 | 5 | 0.21 |
| 0.3 | 0.8 | 5 | 1 | 4 | 0.44 |

**mBAF**: Observed mirrored BAF signal, **BAF**: Observed BAF signal, **CN**: Tumour's Copy Number, $N_A^t$ : Tumour's A allele amount, $N_B^t$ : Tumour's B allele amount, $\alpha$ : % of tumour cells.

**Table 4:** Classical BAF signal confusion example.

| **mBAF** | **BAF** | **CN** | $N_A^t$ | $N_B^t$ | $\alpha$ |
| --- | --- | --- | --- | --- | --- |
| 0.3 | 0.2 | 1 | 1 | 0 | 0.75 |
| 0.3 | 0.2 | 2 | 2 | 0 | 0.6 |
| 0.3 | 0.2 | 3 | 3 | 0 | 0.5 |
| 0.3 | 0.2 | 4 | 4 | 0 | 0.43 |
| 0.3 | 0.2 | 5 | 4 | 1 | 1 |
| 0.3 | 0.2 | 5 | 5 | 0 | 0.38 |
| 0.3 | 0.8 | 1 | 0 | 1 | 0.75 |
| 0.3 | 0.8 | 2 | 0 | 2 | 0.6 |
| 0.3 | 0.8 | 3 | 0 | 3 | 0.5 |
| 0.3 | 0.8 | 4 | 0 | 4 | 0.43 |
| 0.3 | 0.8 | 5 | 0 | 5 | 0.38 |
| 0.3 | 0.8 | 5 | 1 | 4 | 1 |

**mBAF**: Observed mirrored BAF signal, **BAF**: Observed BAF signal, **CN**: Tumour's Copy Number, $N_A^t$ : Tumour's A allele amount, $N_B^t$ : Tumour's B allele amount, $\alpha$ : % of tumour cells.

**Figure 3.**
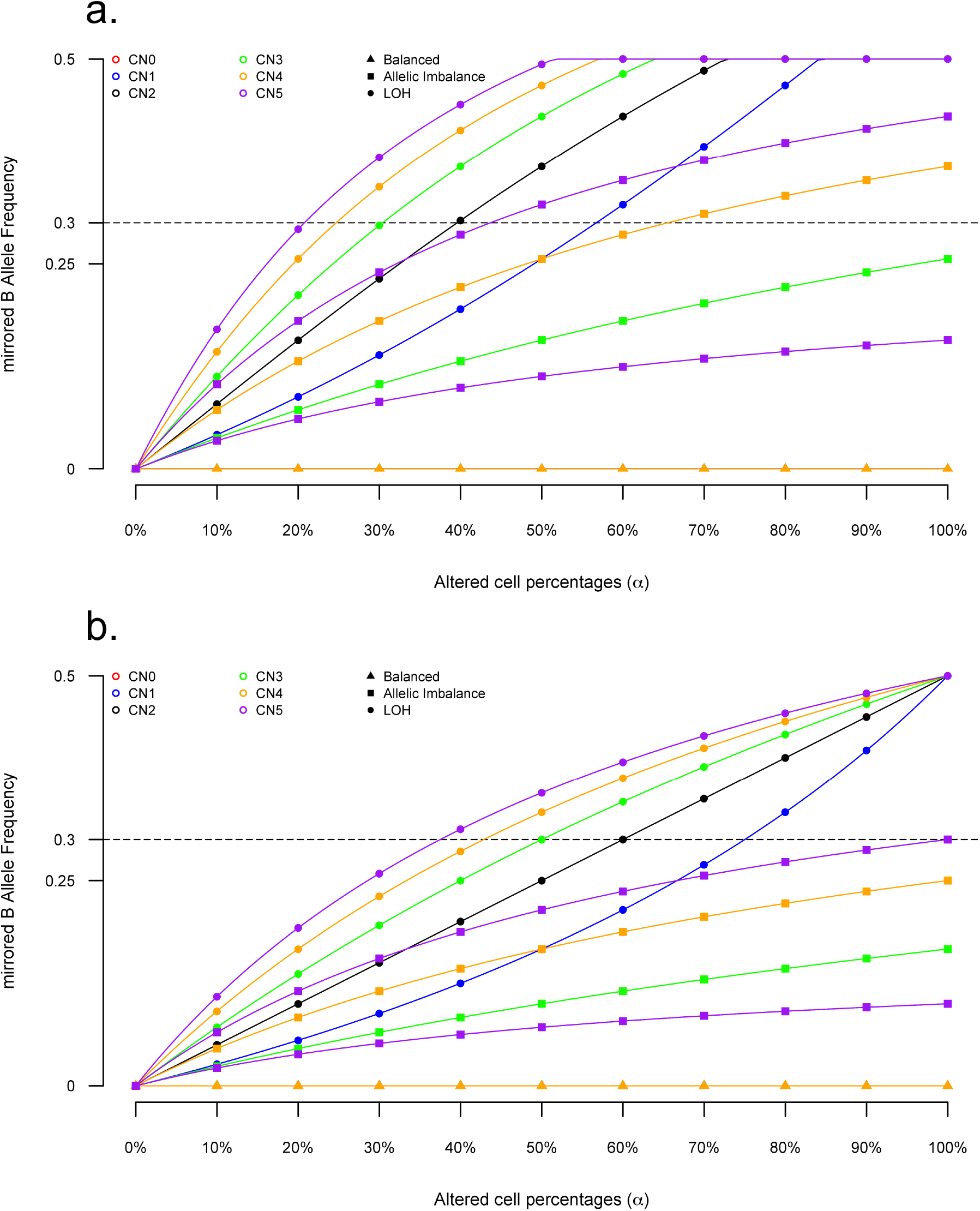
Signal Confusion in mBAF analysis. Evolution of the mBAF signal against α% for CNs from 0 to 5 (red to purple) which result from three genotype classes. Balanced genotypes (triangles). LOH genotypes (circles). Other Ai genotypes (squares). Each value of the graph was simulated using the following algorithms. **a**. Wang’s Θ derived mBAF. **b**. Classical mBAF derived from SiDCoN algorithm. Dashed lines highlight signal confusion for *mBAF*=0.3 and *mBAF*=0.837. Every mBAF signals can be achieved with different allelic combinations at different α% of altered cells.

#### LOH detection

The LOH being a particular case of Ai, we looked for potential signal distortion that would disable the Ai detection. This detection being a boolean method where only signals above 0 are considered, any signals that were close to 0 will result in a confusion. Thus, to assess this, we looked at any Ai signals that were close to 0 between paired samples. With the exception of α≤0.1, numerous signals can be achieved with different CNs. Thus most signals were detectable and different from 0 (data not shown).

### Combined analysis: Copy Number and Allelic Imbalance

To assess the impact of normal cells on the combined analysis, we looked at the signal confusion that occurred simultaneously for both CN and Ai analysis. These analysis encompass the genotype assignment and the combined LRR and BAF analysis. Again, we simulated all the possible allelic combinations (AA, ABB, *etc*.) of CNs from *CN*_0_ to *CN*_5_ diluted with an increasing amount of normal cells. The special case of genome wide association studies (GWAS) will be debated in the discussion.

#### Combined R and Θ analysis (Genotype assignment)

Figure 4 summarises the scatterplot of Θ against R values for all genotype combinations that led to CNs from 0 to 5 contaminated with the increasing gradient of normal cells. Almost all allelic combinations showed a unique and distinct signal. However, signal confusion was still observed for the three following allelic combinations, 1. AAAAB for *CN*_5_, 2. AAAB for *CN*_4_ and 3. AAB for *CN*_3_, as well as their symmetric combinations for the B allele. As exemplified by the arrows and reported in the table 5, a Θ signal of 0.3 combined to a R signal of 2.963 can be achieved with the following α and genotype combinations. 1. α=0.32 (AAAAB), 2. α=0.48 (AAAB), and 3. α=0.96 (AAB). Any R signal can be estimated from a Θ, with the equation 10 (see Methods), obtained based on the equality “α_*R*_ =α_Θ_ “.

**Table 5:**
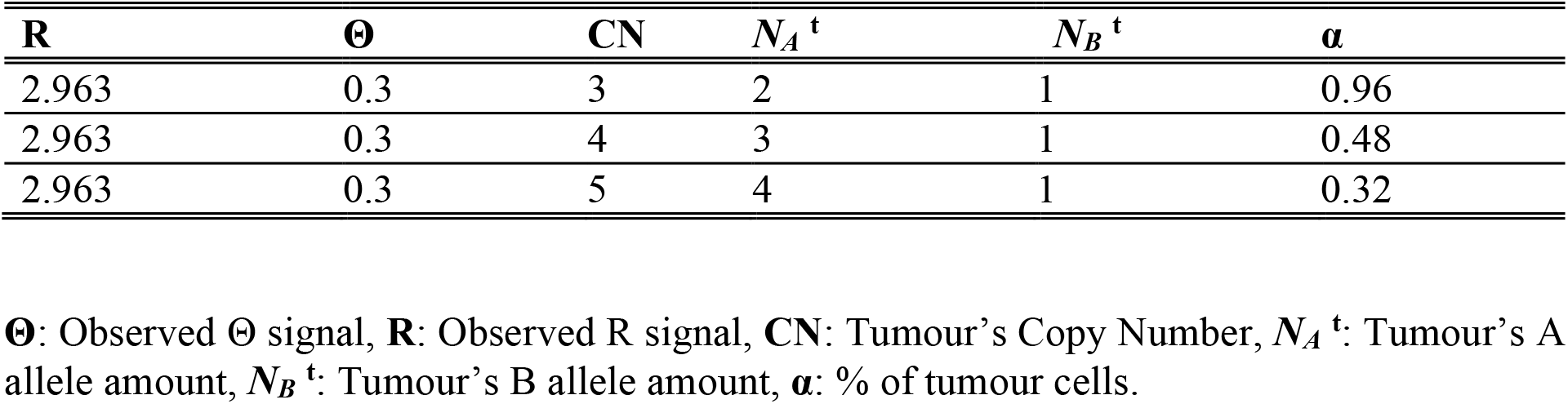
Combined Θ–R’s signal confusion example.

| **R** | **$\Theta$** | **CN** | **$N_A^t$** | **$N_B^t$** | **$\alpha$** |
| --- | --- | --- | --- | --- | --- |
| 2.963 | 0.3 | 3 | 2 | 1 | 0.96 |
| 2.963 | 0.3 | 4 | 3 | 1 | 0.48 |
| 2.963 | 0.3 | 5 | 4 | 1 | 0.32 |

**$\Theta$** : Observed $\Theta$ signal, **R**: Observed R signal, **CN**: Tumour's Copy Number, $N_A^t$ : Tumour's A allele amount, $N_B^t$ : Tumour's B allele amount, **$\alpha$** : % of tumour cells.

**Figure 4.**
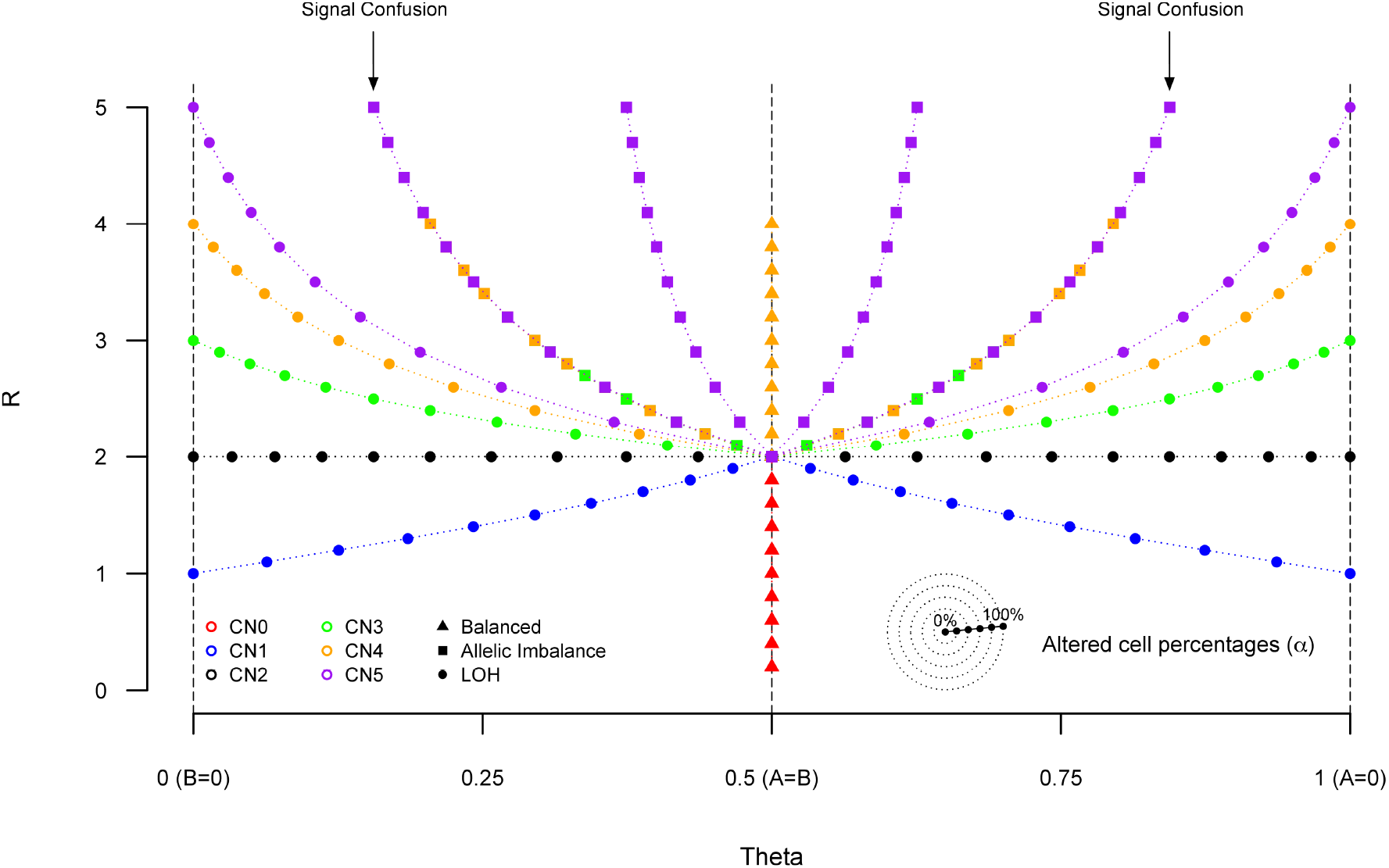
Signal Confusion in genotype assignment. Co-evolution of the R and Θ signals against α% for CNs from 0 to 5 (red to purple), which result from the following genotype classes. Balanced genotypes (triangles). LOH genotypes (circles). Other Ai genotypes (squares). Each Θ value of the graph was simulated using Wang’s method. Each R value of the graph was simulated using the equation *R*_*i*_=*A*_*i*_+*B*_*i*_. Each curve describes an increasing α% of altered cells from the center (Θ=0.5,*R*=2) to the extremes. The two arrows highlight the signal confusion at the genotypes AAAAB for *CN*_5_, AAAB for *CN*_4_ and AAB for *CN*_3_ and their mirror for the B allele. For these genotypes, the same combined *R*−Θ signal can be achieved with different α% of altered cells.

#### Combined LRR and BAF analysis

Following the same principle, we looked for signal confusions that occurred both in the LRR and the BAF signals for each combination of genotype and α. As reported in table 6, a Wang’s BAF signal of 0.4 and a LRR signal of 0.117 can be achieved with the following α and genotype combinations. 1. α=0.1 (AAAAB), 2. α=0.14 (AAAB), and 3. α=0.29 (AAB). Similarly, as reported in table 7, a classical BAF of 0.39 and a LRR signal of 0.216 can be achieved with the following α and genotype combinations. 1. α=0.19 (AAAAB), 2. α=0.28 (AAAB), and 3. α=0.56 (AAB).

**Table 6:** Combined Θ based BAF–LRR’s signal confusion example.

| **LRR** | **mBAF** | **BAF** | **CN** | $N_A^t$ | $N_B^t$ | $\alpha$ |
| --- | --- | --- | --- | --- | --- | --- |
| white 0.117 | 0.1 | 0.4 | 5 | 4 | 1 | 0.1 |
| white 0.117 | 0.1 | 0.4 | 4 | 3 | 1 | 0.14 |
| white 0.117 | 0.1 | 0.4 | 3 | 2 | 1 | 0.29 |
| white 0.117 | 0.1 | 0.6 | 5 | 1 | 4 | 0.1 |
| white 0.117 | 0.1 | 0.6 | 4 | 1 | 3 | 0.14 |
| white 0.117 | 0.1 | 0.6 | 3 | 1 | 2 | 0.29 |

**LRR**: Observed Log R ratio, **CN**: Tumour's Copy Number, $N_A^t$ : Tumour's A allele amount, $N_B^t$ : Tumour's B allele amount, $\alpha$ : % of tumour cells.

**Table 7:** Combined Classical BAF–LRR’s signal confusion example.

| **LRR** | **mBAF** | **BAF** | **CN** | $N_A^t$ | $N_B^t$ | $\alpha$ |
| --- | --- | --- | --- | --- | --- | --- |
| white 0.216 | 0.11 | 0.39 | 5 | 4 | 1 | 0.19 |
| white 0.216 | 0.11 | 0.39 | 4 | 3 | 1 | 0.28 |
| white 0.216 | 0.11 | 0.39 | 3 | 2 | 1 | 0.56 |
| white 0.216 | 0.11 | 0.61 | 5 | 1 | 4 | 0.19 |
| white 0.216 | 0.11 | 0.61 | 4 | 1 | 3 | 0.28 |
| white 0.216 | 0.11 | 0.61 | 3 | 1 | 2 | 0.56 |

**LRR**: Observed Log R ratio, **CN**: Tumour's Copy Number, $N_A^t$ : Tumour's A allele amount, $N_B^t$ : Tumour's B allele amount, $\alpha$ : % of tumour cells.

### Signal confusion estimation

Based on the signal confusions detected within the previous analysis, we evaluated the probability of a signal confusion occurrence in the classical SNP–aCGH analysis.

#### LRR signal confusion

Figure 5 summarises the probability of a confusion occurrence during the analysis of a candidate LRR signal for cells that carry up to 5 chromosomal copies under an expected *CN*_*max*_ from 0 to 10. Interestingly, the signal confusion probability remains exactly the same along the expected *CN*_*max*_ for the *CN*_0_ :*p*_*c*_ =0, *CN*_1_ :*p*_*c*_ =0.5 and *CN*_2_ :*p*_*c*_ =0. Otherwise, similar shapes were observed for gains. In this context, the probability is higher for lower CN gains when assessed with any expected *CN*_*max*_. Thus, the probability to get a confused signal with an expected *CN*_*max*_ =10, is higher for the *CN*_3_ than any other upper CNs.

**Figure 5.**
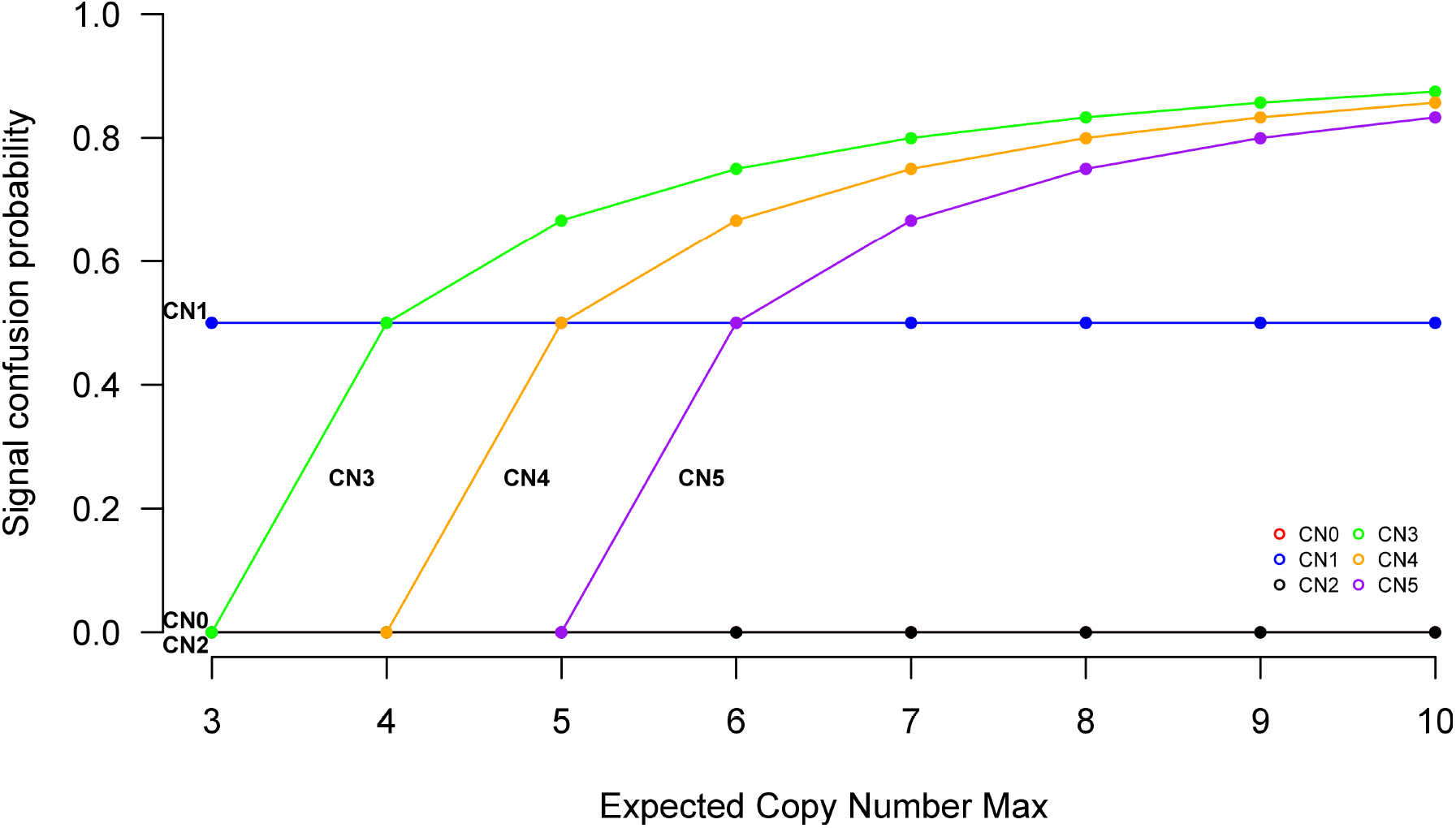
Signal confusion occurrence in LRR signal. Evolution of the signal confusion probability against the expected *CN*_*max*_ of tumour cells that carry CNs from 0 to 5 (red to purple). Each value of the graph was obtained using the equation 11 (See Methods). Deleted (*CN*_0_, *CN*_1_) and normal (*CN*_2_) copy numbers are *CN*_*max*_ independant. Gains (*CN*>2) describe the same curve shape, which proportionally increases along with the *CN*_*max*_ expectation. Lower gains lead to a high signal confusion.

#### Θ signal confusion

Figure 6 summarises the probability of a confusion occurrence during the analysis of a candidate genotype’s Θ signal for cells that carry up to 5 chromosomal copies under an expected *CN*_*max*_ from 0 to 10. Interestingly, the signal confusion probability reaches exactly the same values for any LOH under the same *CN*_*max*_. For oAis, however, the probabilities follow independent curves. Furthermore, the more unbalanced the genotype, the lower the probability is.

**Figure 6.**
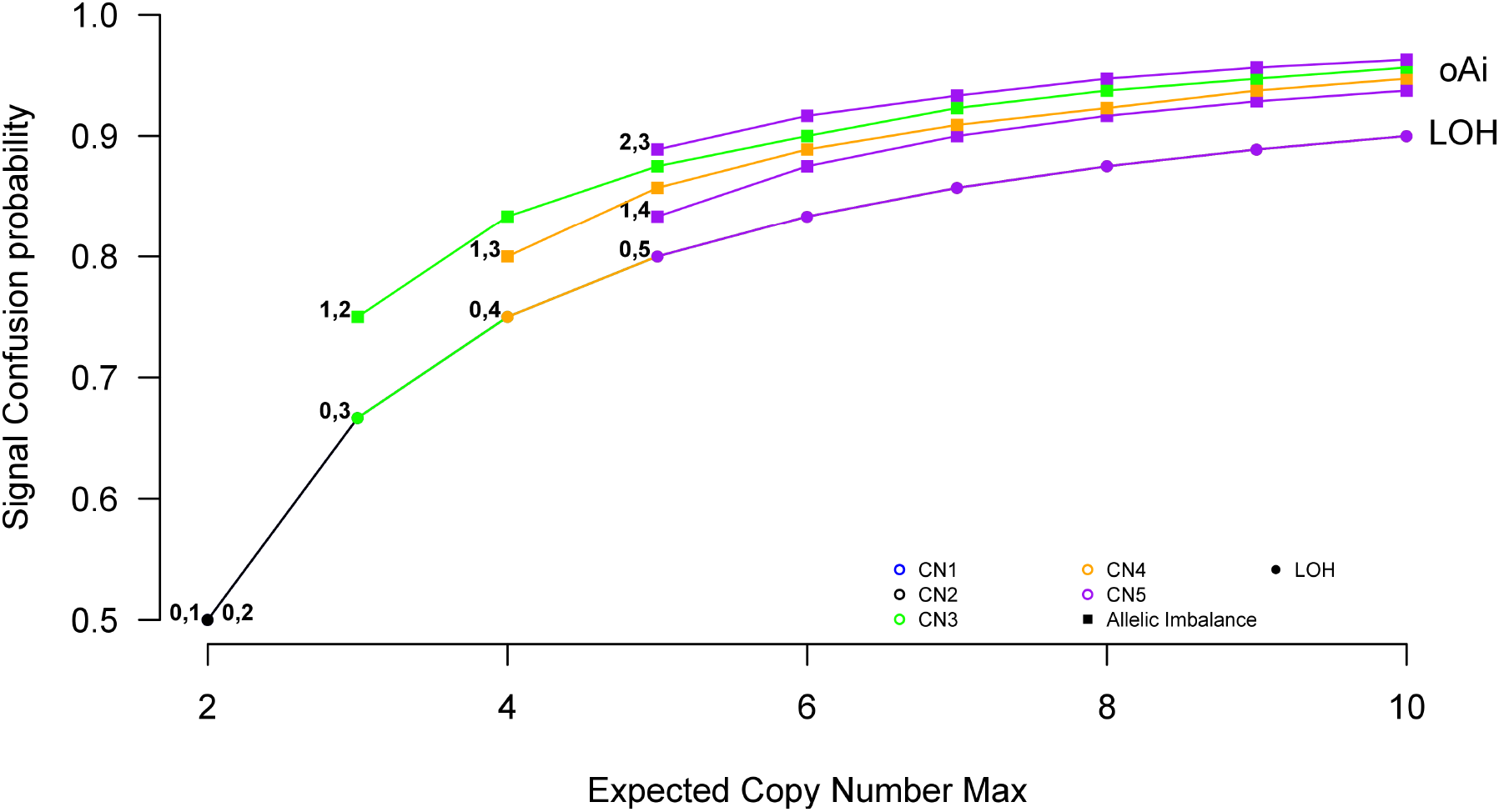
Signal confusion occurrence in theta signal. Evolution of the signal confusion probability against the expected *CN*_*max*_ of tumour cells the possible genotypes for CNs from 1 to 5 (blue to purple) which result from two genotype classes: LOH genotypes (circles). Other Ai genotypes (squares). LOH: Loss of Heterozygosity. oAi: Other Allelic imbalance. Each value of the graph was obtained using the equation 16 (See Methods). LOHs share a common curve shape and are the lowest probable confusion for any expected *CN*_*max*_. oAi describe independent curve with the same shape, which increases along with an increasing *CN*_*max*_ expectation. The most unbalanced genotypes show the lowest probability of signal confusion.

## Discussion

In an effort to elucidate the impact of the tumour contamination with stromal cells on classical SNP–aCGH analysis, we simulated genotype data that led to copy number aberrations from *CN*_0_ to *CN*_5_ diluted by an increasing gradient of normal cells. With only one tumour population contaminated by normal cells, we showed that signal confusion can disturb the interpretation of each classical SNP-aCGH analysis.

Regarding the copy number analysis, LRR signals were shown with a substantial signal confusion for both deletions and gains. For deletions (figure 1 bottom) all simulated *CN*_1_ values can be achieved with *CN*_0_ values at another α% of tumour cells. For gains (figure 1 top) the confusion was dependant on the CN maximum (*CN*_*max*_). In our simulations, *CN*_*max*_ was set to 5, and thus the confusion encompassed up to three CNs. While previous studies considered a *CN*_*max*_ up to 4, we demonstrated that higher CNs may have a dramatic impact on the data interpretation. However, higher CNs can not be excluded when the exact CN have to be assessed, especially in the double minute context [25, 26]. As shown by our results, this assessment, based only on the LRR analysis, was highly similar to the problem already mentioned for mixtures of different tumour populations [4]. In this context, we expect signals from oligo or BAC array CGH (aCGH) to be strongly biased by these confusions when analysing contaminated samples. Indeed, these platforms do not allow the allele specific analysis. Nevertheless, the probability to have a confusion is higher for the gains than the deletions (figure 5). Indeed, among the deletions, only the *CN*_1_ can be misinterpreted during the CN assignment, and this for any expected *CN*_*max*_. Contrariwise, the confusion for a gain is even more probable while the expected *CN*_*max*_ is high. Importantly, the lower the observed signal is, the highly confused the assignment might be.

At the Allelic imbalance level, while the Θ based BAF enabled a better separation of the signals, Θ only or BAF analysis were shown with the highest signal confusion among the SNP-aCGH analysis. Interestingly, while each class was the subject of different signal confusions, the Θ based BAF showed more confusions than the classical BAF (one more genotype combination can reach the signal BAF=0.3). As displayed by figures 2 and 3, three classes of genotypes can be the source of the same signals. 1. Balanced (triangles). 2. LOH (circles). 3. Other Ais (squares). Every signals coming from classes 2 can be achieved with all other possible genotypes. Only one signal was observed for every genotype of the class 1. Signals coming from class 3 are more confused when the genotype tends to be more balanced. As these three classes were equally probable when deriving from an heterozygous genotype, the expected signal is thus more confused while it tends to a balanced genotype acquired by the tumour (figure 6). Interestingly, any kind of pure LOH are equally probable, when available under the same *CN*_*max*_ expectation. Besides, all aberrant genotypes that derived from the homozygous genotypes (AA or BB), led to the same constant signal of LOH, thus to a constant confusion. Consequently, algorithms that are mainly based on Ai signals can fall into a signal confusion, especially when trying to discover the exact percentage of normal and tumour cells in the sample. Nevertheless, the LOH detection analysis should not suffer from this confusion, as no exact genotype have to be assessed.

The combination of both CN and Ai analysis was widely used in previous studies [9, 10, 14, 19, 21, 22]. Most often, it was mentioned in these studies that the combined CN and Ai analysis can help with the normal cell contamination. Indeed, it can remarkably increase the power of the assessment both for the exact CN_*max*_ and the normal cell contamination. However, our results showed that this combined analysis can also led to a signal confusion. As shown by the figure 4, we highlighted three genotypes, AAAAB, AAAB and AAB, which displayed similar signals. As *CN*_*max*_ was set up to 5 for the present study, we expect higher CNs to results in even more confusion. Moreover, it is worth to note that the signal confusion should be even more important for low α under high background noise conditions (figure 4, center). Nevertheless, in the context of the current simulations, only one tumour population was studied without any background noise. In this context, the observed confusions should be solved using a pigeon hole logic based on the whole set of observed aberrations. For more complex tumour cases, however, the analysis can turn into an ill-mathematical problem [4]. The problem falls therefore under Parisi’s conclusions, where more than 3 populations within the same sample result in numerous issues, even if a pigeon hole logic is used. Importantly, as a consequence of this signal confusion that may occur during the genotyping step, any subsequent tumour genome-wide association studies (GWAS) would also suffer from the normal cell contamination. As demonstrated in figure 4, three cases could be interpreted as one unique genotype. However, common genotype calling algorithms can only, somehow, average these combinations, thus flattening the true underlying genotype [27, 28]. Beside this unavoidable confusion, classical statistical approaches are subject to a loss of power as the number of degrees of freedom increase with additional genotypes resulting from the various alterations commonly found in tumours. Under such conditions, GWAS are approximated, thus missing numerous potential associations.

Studies that suggested the use of a three copy–number state (gain, deletion, normal), can circumvent the exact copy–number assessment, but will loss the tumour content information [4]. Because this exact copy number status couldn’t be easily assessed for a punctual analysis, a pigeon hole logic analysis is often used. However, these algorithms have to make a choice between multiple scenario of numerous individual alterations. The signal confusion probabilities, developed in the current study, weight each alteration and let the user aware about all other possible scenario that were omitted. For example, in the simple case of 5 deletions, 3 *CN*_1_ and 2 *CN*_0_, the chosen scenario will be one among height others. With ***p***_***u***_^***CN1***^ ***– 0*.*5*** and ***p***_***u***_^***CN0***^ ***– 1*** the scenario probability is **0.5**^**3**^ **\* 1**^**2**^ **– 1/8**. Consequently, the association between the developed equations (11, 16 and 17) and a pigeon hole logic can greatly help the diagnostic decision about the robustness of the scenario chosen during the exact copy number assignment.

## Conclusions

Early identification of tumour populations that recur after current therapies is crucial for future diagnosis, prognosis and patient’s care [3, 5, 29, 30]. Studies that attempted to unveil the tumour content, were important steps to build robust diagnosis tools [8, 25, 31]. However, tumour samples remain complex mixtures with an unknown number of populations, including a random amount of normal cells. In this context, an unknown number of issues are possible in the tumour analysis [4, 9]. While we accepted the relevance of previously discovered tumour markers, omitting the signal confusion coming with the stromal contamination would lead to an incorrect assessment of the tumour content. Indeed, we showed here that the normal contamination can itself lead to numerous interpretation of the tumour alterations. Moreover, the signal confusion remained present in all kind of CGH microarray analysis. Being hard to circumvent, nevertheless, this confusion can be assessed. Only few genotypes were actually the source of a constant confusion in every analysis. Therefore, the current work proposed a way to appreciate how many other possible scenario would lead to the same altered signals. Rather than a direct assessment of the exact copy–number, we proposed here a degree of certitude regarding a signal confusion occurrence. Combining these degrees of certitude for each alteration with a pigeon hole logic can improve the robustness of the tumour sub-populations identification. Such probabilities could also be integrated in a three state analysis (gains, deletions and normal) to help clinicians in their decisions. Understanding how the normal contamination acts on the tumour informations, can really make a difference in the assessment of the tumour content and thus in the early identification of a treatment response. Subsequent analysis of sub-clonal impact on drug response, are likely to have a major impact on the development of new treatment strategies and patient’s care.

## Methods

### Reference and sample’s genotypes

Thousand reference samples were generated with the following genotype proportions, 35% of AA, 35% of BB and 30% of AB. Based on these genotypes, all possible combinations of tumour genotypes were generated according to the type of CN evaluated. For example, for a *CN*_3_, the genotypes AAA, AAB, ABB and BBB were evaluated if the corresponding reference genotype was AB. In the same context, The genotype AAA was evaluated if the corresponding reference genotype was AA.

### Tumour and Reference proportions

An increasing gradient of altered tumour cells (α) was used to simulate dilution of tumour cells with normal cells. Briefly, each analysis was weighted with α. The tumour signal (*S*_*t*_) weighted by α (*Sw*_*t*_), was obtained with *Sw*_*t*_ =α.*S*_*t*_. Similarly, normal cell signal (*S*_*n*_) weighted by α (*Sw*_*n*_), was obtained with *Sw*_*n*_ =(1−α).*S*_*n*_.

### Allele intensities

For each simulated *i*^*th*^ SNP probe, weighted *A*_*i*_ and *B*_*i*_ allelic intensities were obtained based on the allele proportions in the tumour (***N***_***X***_^**t**^) and in the reference (***N***_***X***_^***n***^) genotypes, such as

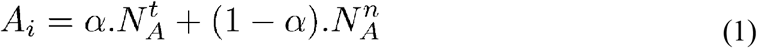

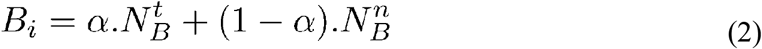

### Copy Number analysis

Weighted DNA quantities, for each *i*^*th*^ SNP probe (*R*_*i*_), were obtained by summing the weighted allele intensities (equations 1 and 2), such as *R*_*i*_ =*A*_*i*_ +*B*_*i*_. The subsequent R ratio (*RR*_*i*_) and Log R ratio (*RR*_*i*_) were obtained based on SiDCoN method with *RR*_*i*_ =*R*_*i*_ /2 and *LRR*_*i*_ =*log*_10_ (*RR*_*i*_).

### Allelic analysis

#### Theta Θ

Weighted Θ values, for each *i*^*th*^ SNP probe (Θ_*i*_), were obtained based on weighted allele intensities (equations 1 and 2), using Wang *et al* methodology [24], such as

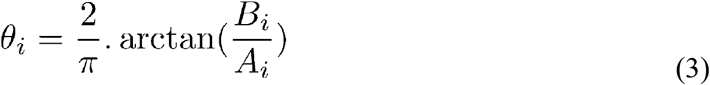

#### BAF and mBAF

Two distinct methods were compared to simulate weighted BAF values for each *i*^*th*^ SNP probe (*BAF*_*i*_). The first method was based on Θ values and was previously described by Wang *et al* [24]. The second method was described with the SiDCoN algorithm [23], and was used here as follow

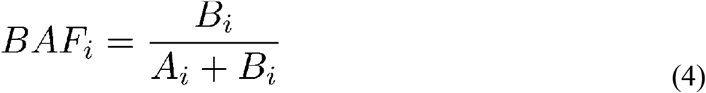

The weighted mirrored BAF signal for each *i*^*th*^ SNP probe (*mBAF*_*i*_) was obtained for both methods using the following equation

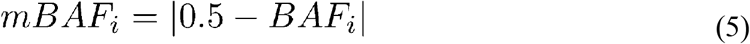

### LOH detection analysis

Following the same principle, weighted Ais were used to assess if the signal confusion can disturb the Ai and subsequent LOH detection. This detection is boolean method where only the difference in allele frequency between the tumour and the paired constitutive reference samples is assessed. We used previously published method called | dAlleleFreq *tum*−*ref*|, the absolute value of this allele frequency difference [20, 32]. Since both samples are from the same patient, this allele frequency difference is closed to zero except in regions exhibiting Ais or LOHs. Possible confusions were therefore detected when any Ai or LOH signals were closed to zero.

### Linear models

#### LRR analysis

Any α from the figure 1 can be inferred based on the *LRR* signal, using the following equation.

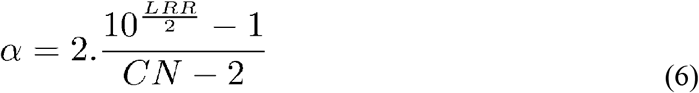

Importantly, equation 6 returned non interpretable α for *LRR*<0 and *CN*>2, *LRR*>0 and *CN*<2 or any LRR and CN = 2, this last case always returned aberrant results as any α values between 0 and 1 are possible for the *CN*_2_.

#### Theta (Θ) analysis

Any α from the figure 2 (a) can be inferred based on the Θ signal using the following equation.

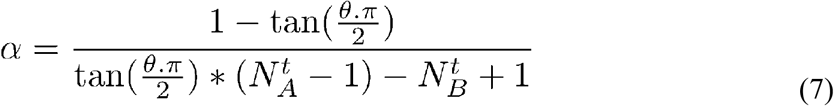

Where ***N***_***A***_ ^**t**^ and ***N***_***B***_ **^t^** are the amount of A and B of the considered genotype. Importantly, equation 7 returned non interpretable α for Θ<0.5 and ***N***_***A***_ ^**t**^ **< *N***_***B***_ **^t^**, Θ>0.5 and ***N***_***A***_ ^**t**^ **> *N***_***B***_ ^**t**^ or any Θ and ***N***_***A***_ ^**t**^ **- *N***_***B***_ **^t^**, this last case always returned aberrant results as any α values between 0 and 1 are possible and Θ will always be equal to 0 for balanced genotypes.

#### BAF analysis

Any α from the figure 3 (a) can be inferred from the Θ based BAF signal, using the following system of equations 8 to obtain Θ and then using the previous equation 7 to obtain α.

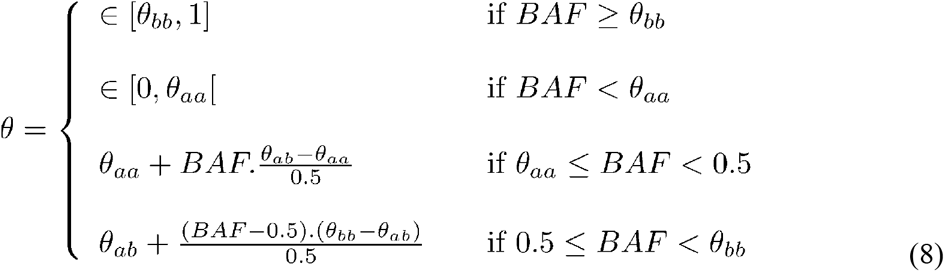

Besides, any α% from the figure 3 (b) can be inferred from the classical BAF signal, using the following equation.

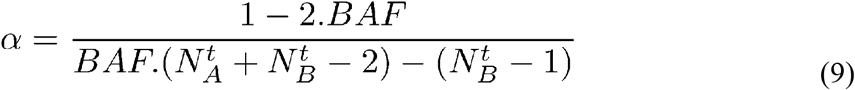

Where ***N***_***A***_ ^**t**^ and ***N***_***B***_^**t**^ are the amount of A and B alleles respectively. Importantly, this equation 9 returned non interpretable α for the following combinations. 1. *BAF*<0.5 and ***N***_***A***_^**t**^ **< *N***_***B***_ ^**t**^, 2. *BAF*>0.5 and ***N***_***A***_ ^**t**^ **> *N*_*B*_ ^t^** or 3. any *BAF* and ***N***_***A***_^**t**^ **- *N***_***B***_ ^**t**^, this last case always returned aberrant results because any α values between 0 and 1 are possible and thus the BAF will always be equal to 0 for balanced genotypes.

#### Genotype analysis

Any *R* values from the figure 4 can be retrieved from the Θ value using the α equality 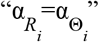, which result in the following equation.

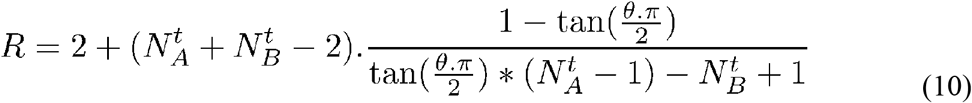

### Signal confusion, interference estimation

#### Interference in LRR

The probability ***p***_***c***_ ^***LRR***^ to measure a confused LRR signal is **1 − *p***_***u***_^***LRR***^, probability of measuring a unique LRR signal. ***p***_***u***_^***LRR***^ was obtained in two steps. 1. A classical Hidden Markov Model was used to discretise the observed signal *S*_*o*_ into CN status (CN). 2. The probability was estimated based on the following system of equations.

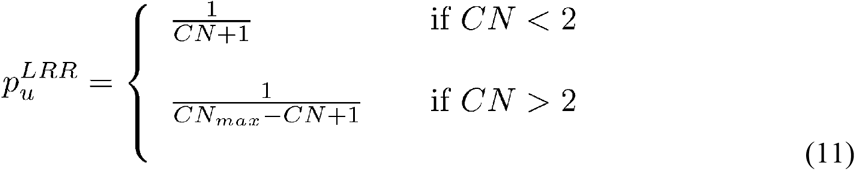

Where *CN*_*max*_ is the expected maximal CN status for the sample, which had to be fixed based on expectation.

#### Interference in Θ

The probability ***p***_***c***_^Θ^ to measure a confused Θ value is **1- *p***_***u***_^Θ^, probability of measuring a unique Θ value. This ***p***_***u***_^Θ^ was measured for Θ∈[0.5,1]. Since Θ values can be divided in three distinct signal shapes, three distinct ***p***_***u***_^Θ^ were assessed, balanced and allelic imbalanced (LOH and oAi) genotypes. In all three cases, the ***p***_***u***_^Θ^ depends on the expected CN maximum (*CN*_*max*_).

1. ***p***_***u***_^ΘLOH^ for LOH genotypes, achieved for Θ ∈ (0.5,1].
2. ***P***_***u***_ ^ΘBAL^ for Balanced genotypes, achieved for Θ_*obs*_ =0.5
3. ***p***_***u***_^ΘoAi^ for other Ai genotypes, achieved for for Θ ∈ (0.5,1).

1. Since only one LOH genotype is expected per CN, any Θ_*obs*_ value can be achieved with *CN*_*max*_ genotypes. Thus the associated probability is defined by:

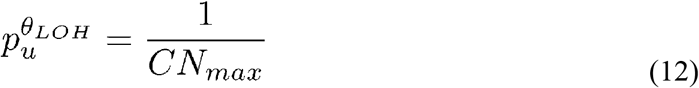
2. As only even CNs can reach a Balanced genotype, the value of Θ_*obs*_ =0.5 can be achieved with 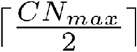 genotypes. Thus the associated probability is defined by:

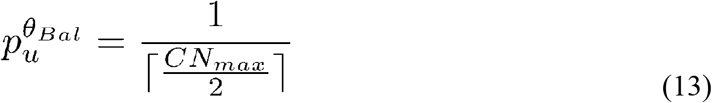
3. To assess the probability among oAi signals, we first discretise the theoretical values of every pure genotype signals. These values were obtained with equation 3 for α=1, and were then turned into z levels, with z ∈ [1, zmax] and *z*_*max*_ defined with the following equation.

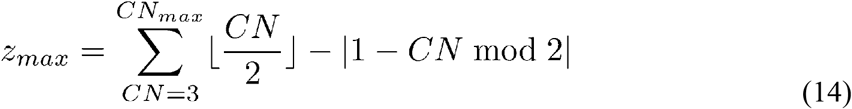

Thus, an observed Θ value will fall in Θ_*obs*_ ∈ [z, z+1), and the ***p***_***u***_^ΘoAi^ is obtained with the following equation:

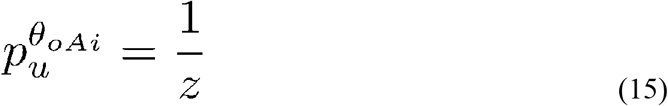

Combining the previous equations 12 and 15, ***p***_***u***_^ΘAi^ was obtained for Θ_*obs*_>0.5 with the following equation.

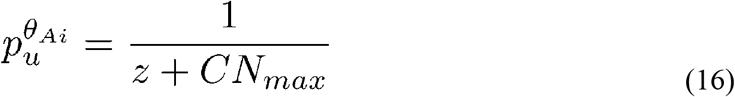

In this context, z levels were assigned based on the previous method, *z*_*max*_ +1 for Balanced genotypes and *z*=0 for LOH genotypes. All theoretical values for z levels can be retrieved using the equations 1, 2 and 3 with α=1 (see Methods).

In the example of figure 2, the expected maximal CN is *CN*_*max*_ =5, which lead to *z*_*max*_ =4. The Θ_*obs*_ =0.837 belongs to Θ_*obs*_ ∈ [z=1, z+1=2). A strong probability for a confused signal is therefore obtained using equation 16, such as ***p***_***c***_^Θ^ - 1 – ***p***_***u***_^Θ^ - 1 – (1/(1+5)) – 0.834.

#### Interference in combined *R*−Θ

Most of the combined *R*−Θ values were unique (figure 4). The signal confusion appeared only for three genotypes, AAAAB, AAAB and AAB as well as their mirror for the B allele. Consequently, the signal confusion probability was defined only for these specific cases with the following equation.

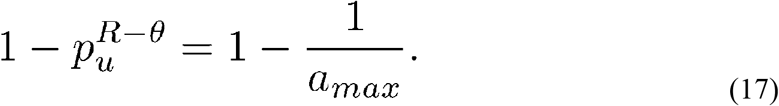

where, *a*_*max*_ is the maximum amount of A allele obtained for a balanced genotype within a maximal CN (*CN*_*max*_) and follows *a*_*max*_ – [*CN*_max_ / 2].

